# KMT5C displays robust retention and liquid-like behavior in phase separated heterochromatin

**DOI:** 10.1101/776625

**Authors:** Hilmar Strickfaden, Kristal Missiaen, Michael J. Hendzel, D. Alan Underhill

## Abstract

The pericentromere exists as a distinct chromatin compartment that is thought to form by a process of phase separation. This reflects the ability of the heterochromatin protein CBX5 (*aka* HP1*α*) to form liquid condensates that encapsulate pericentromeres.^1,2^ In general, phase separation compartmentalizes specific activities within the cell, but unlike membrane-bound organelles, their contents rapidly exchange with their surroundings.^3^ Here, we describe a novel state for the lysine methyltransferase KMT5C where it diffuses within condensates of pericentromeric heterochromatin but undergoes strikingly limited nucleoplasmic exchange, revealing a barrier to exit similar to that of biological membranes. This liquid-like behavior maps to a discrete protein segment with a small number of conserved sequence features and containing separable determinants for localization and retention that cooperate to confer strict spatial control. Accordingly, loss of KMT5C retention led to aberrant spreading of its catalytic product (H4K20me3) throughout the nucleus. We further found that KMT5C retention was reversible in response to chromatin state, which differed markedly for CBX5 and the methyl-CpG binding protein MeCP2, revealing considerable plasticity in the control of these phase separated assemblies. Our results establish that KMT5C represents a precedent in the biological phase separation^4^ continuum that confers robust spatial constraint of a protein and its catalytic activity without progression to a gel or solid.

---

Analyses of the prototypic heterochromatin protein CBX5 indicated that phase separation plays an important role in partitioning of pericentromeric heterochromatin.^1,2^ This is particularly evident in mouse interphase nuclei where pericentromeres from multiple chromosomes form chromocenters.^5^ Despite the stable appearance of such assemblies, they are highly dynamic and their protein constituents rapidly exchange with their surroundings, reflecting low affinity, multivalent interactions that allow proteins to coalesce into distinct aqueous compartments.^6,7^ In this context, CBX5 enrichment within pericentromeric heterochromatin requires binding to trimethyllysine-9 on histone H3 (H3K9me3), which is placed by the lysine methyltransferases SUV39H1 and 2.^8^ CBX5 then recruits KMT5C protein to catalyze histone H4 lysine-20 trimethylation (H4K20me3). In fluorescence recovery after photobleaching (FRAP), CBX5 rapidly exchanged between the pericentromere and nucleoplasm,^9^ whereas SUV39H2 and KMT5C appeared to be immobile when entire chromocenters were bleached.^10^ Although concluding that the latter two proteins created a stable scaffold,^10^ these studies did not assess mobility within chromocenters, which would instead suggest retention within a liquid compartment. To test this fundamentally different model, we queried KMT5C dynamics within mouse chromocenters in comparison to CBX5 and MeCP2, which also exhibits pericentromeric enrichment and nucleoplasmic exchange.^10,11^ As previously shown with full chromocenter bleaching,^10,12^ KMT5C displayed long recovery times, indicating minimal exchange with the surrounding nucleoplasm, and was considerably slower than either CBX5 or MeCP2 (Fig. 1a, Extended Data Videos 1-3). Upon partial bleaching, however, KMT5C fluorescence recovered (Fig. 1a, Extended Data Videos 4 and 5) and progressed from the non-bleached portion of the chromocenter (Fig. 1b). This occurred on a timescale where there was no appreciable recovery of fully bleached chromocenters in the same nucleus (Fig. 1b), establishing that KMT5C moved readily within chromocenters but did not efficiently exchange (Fig. 1c). In the context of phase separation, this signifies a remarkable barrier to exit that has not been observed for other proteins without transition to a solid or gel state, which renders them immobile.^13^ KMT5C therefore demonstrates that phase separation can achieve robust compartmentalization while nevertheless retaining liquid-like mobility.

**Figure 1.**
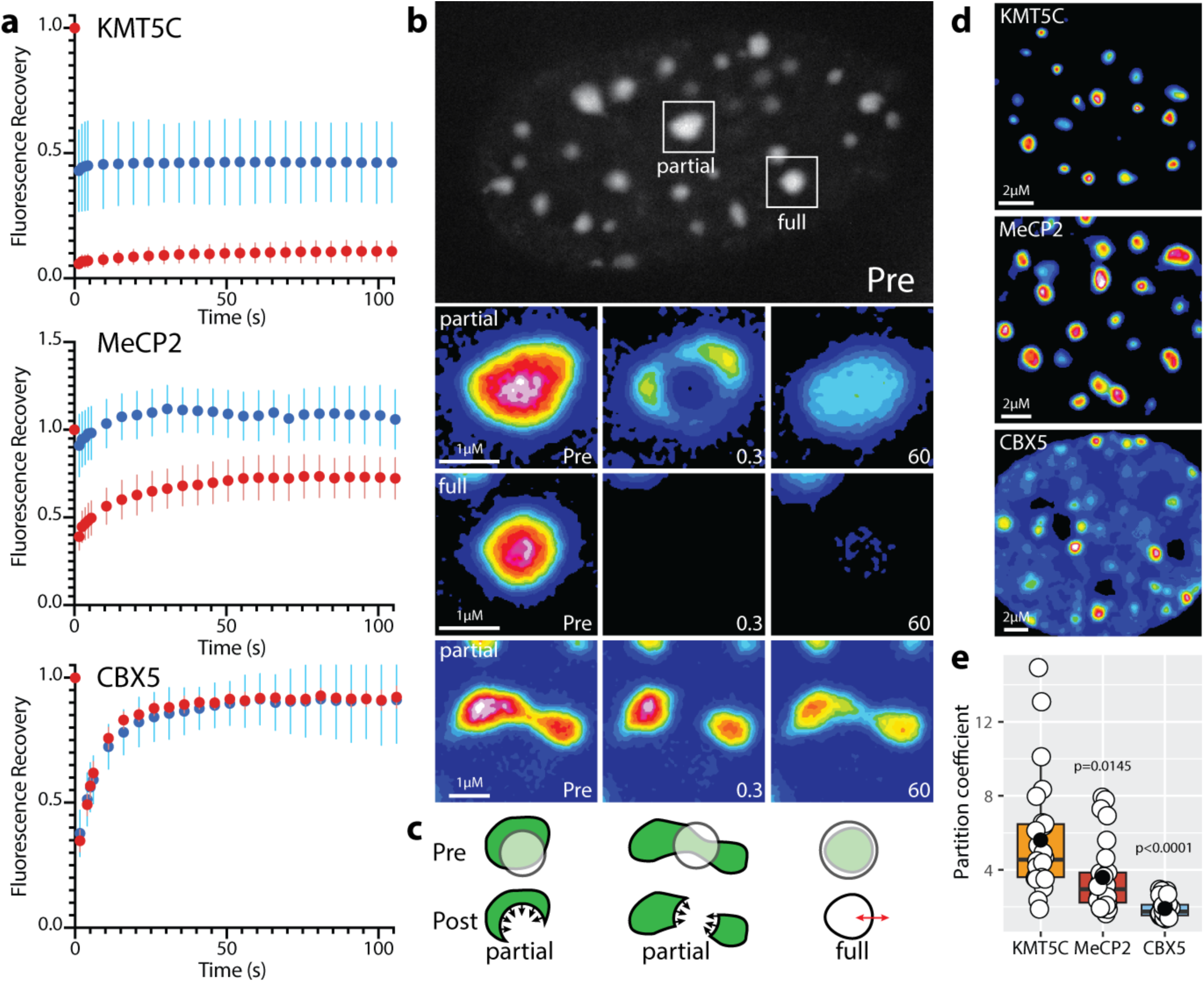
KMT5C is mobile within chromocenters but undergoes limited nucleoplasmic exchange. **a**, FRAP curves for KMT5C, MeCP2, and CBX5-mEmerald fusion proteins in mouse NMuMG breast cancer cells (*n=30*) (see Methods). Partial (*red*) and full (*blue*) fluorescence recovery curves (*filled circles* represent mean fluorescence intensity, while *vertical lines* indicate standard deviation. **b**, Time-lapse series of *full* and *partial* KMT5C-mEmerald photobleaching (Extended Data Movie 4). Insets use a 16-color intensity map to depict fluorescence recovery upon partial or full bleaching from 0.3 to 60s. The second *partial* panel depicts an independent recovery event involving a chromatin bridge between two chromocenters (Extended Data Movie 5), indicating they form a contiguous phase separated environment. **c**, Schematic representation of KMT5C (*green*) movement (*arrows*) in each of the full and partial bleach (*translucent circles*) time series from panel ***b*. d**, Partition images (16-color intensity map) for KMT5C, CBX5 and MePC2. **e**, Scatter plots display corresponding partition coefficients for KMT5B (*n=26*), MeCP2 (*n=30*), and CBX5 (*n=30*) (see Methods).

Consistent with this behavior, KMT5C was more efficient in chromocenter localization than CBX5 (Fig. 1d, e). Unexpectedly, this enrichment did not simply reflect retention because the unrelated MeCP2 protein had a high partition coefficient (Fig. 1d, e) but underwent nucleoplasmic exchange (Fig. 1a). MeCP2 therefore defined a third state with regard to partitioning and mobility, indicating these parameters were separable and occur across a continuum. Intrinsically disordered regions are also typical of phase-separated proteins,^13,14^ including CBX5.^1,2^ Although these features were highly conserved in disorder plots of CBX5 and MeCP2 from representative mammals, they varied for KMT5C along with charge properties (Fig. 2a, Extended Data Table 1). Whereas MeCP2 had the highest overall percentage of disorder (76.7), KMT5C exhibited the lowest (33.6), lacked extended regions of disorder, and its profile was most divergent (Fig. 2a, Extended Data Table 1). This suggests that while CBX5 and MeCP2 have prototypic features of phase separating proteins, KMT5C achieves demixing through other means. To this end, CBX5 and MeCP2 contain highly conserved domains that confer localization to constitutive heterochromatin (Fig. 2a),^15-17^ which for KMT5C involves a C-terminal region^12^ that we will refer to as the <u>C</u>hromocenter <u>R</u>etention <u>D</u>omain (CRD). Sequence identity within the CRD, however, was limited to 18 of 59 residues across mammals (Fig. 2b) and its disorder potential and charge properties (Fig. 2c, Extended Data Table 1) varied considerably (Fig. 2d). To broadly establish functional relevance of the CRD, we assessed versions from *Homo sapiens* (*Hs*) and *Mus musculus* (*Mm*), together with *Cavia porcellus* (*Cp*) and *Bubalus bubalis* (*Bb*) because they exhibited distinct disorder profiles and the greatest range in pI (8.94-11.92) (Fig. 2c, d). Nevertheless, each CRD derivative displayed robust chromocenter partitioning (Fig. 2e, f) that was not significantly different from full-length KMT5C, together with intra-chromocenter mobility and limited exchange (Fig. 2g, Extended Data Videos 6-9). As a result, the dynamic properties of full-length KMT5C can be entirely recapitulated by this short protein segment. Although best exemplified by the mouse CRD, the behavior was shared by all orthologs, suggesting it is driven by common sequence features that are modulated by species-specific differences.

**Figure 2.**
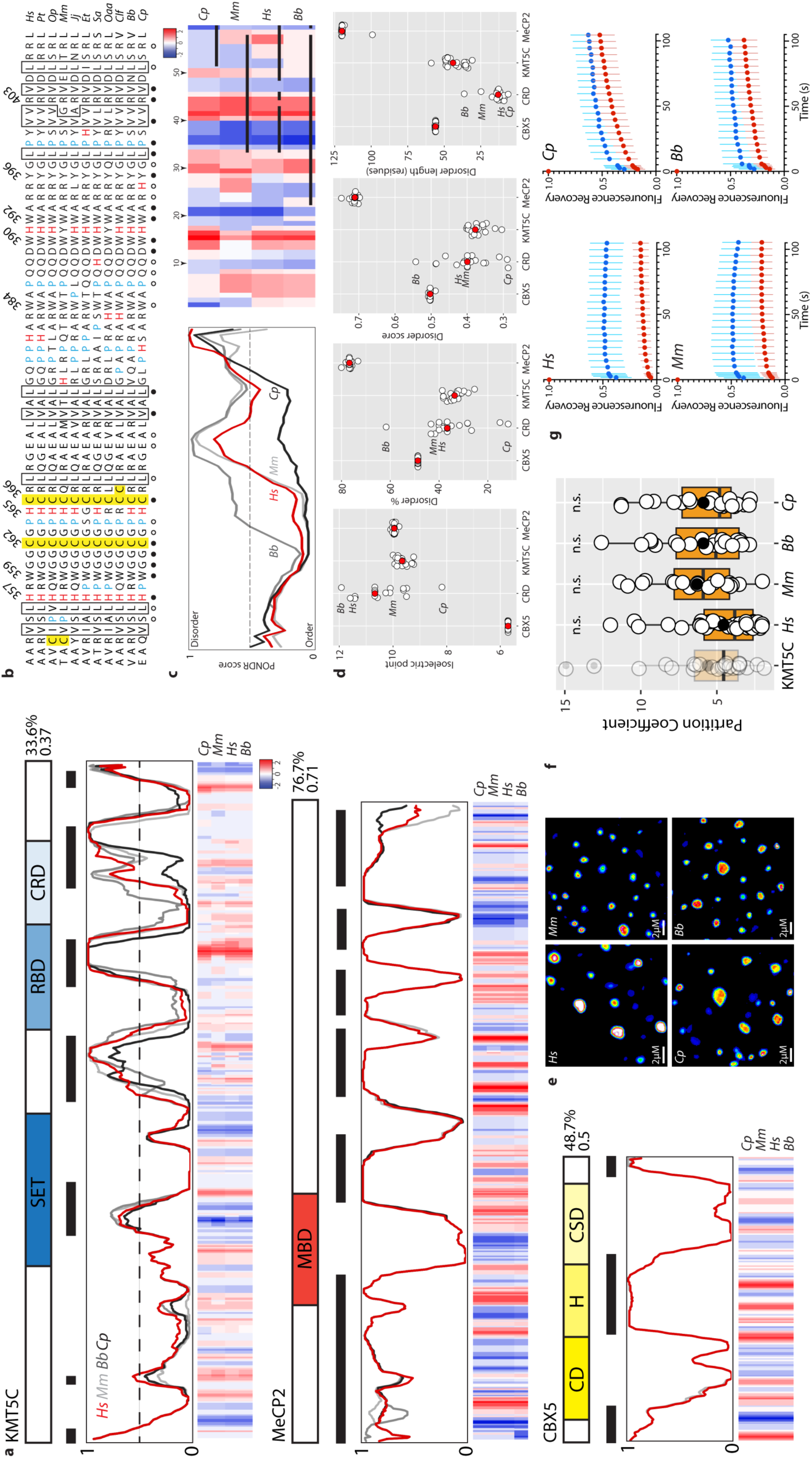
The chromocenter retention domain constrains KMT5C to individual chromocenters. **a**, Disorder plots (PONDR) and charge heatmaps for KMT5C, CBX5, and MeCP2 orthologs from *Homo Sapiens* (*Hs*), *Mus musculus* (*Mm*), *Bubalus bubalis* (*Bb*), and *Cavia porcellus* (*Cp*). For reference, schematics illustrate the distribution of annotated domains in each protein (**S**u(var)3-9, **E**nhancer-of-zeste and **T**rithorax (SET); **R**NA-**B**inding **D**omain (RBD); **C**hromocenter **R**etention **D**omain; **M**ethyl-CpG **B**inding **D**omain (MBD); **C**hromo**d**omain (CD); **H**inge domain (H); and **C**hromoshadow **D**omain (CD)). Input sequences were aligned using Clustal Omega^32^ together with manual removal of gaps and then analyzed for disorder using PONDR^33^ and charge properties using EMBOSS. Average disorder percentage and score are indicated to the *right* of the primary structure schematic, and disordered segments are shown as thick horizontal lines. **b**, Multispecies sequence alignment of the CRD is shown in JPred^34^ format. Residues absolutely conserved in mammals are indicated by a *filled circle*, while those exhibiting 90% conservation are noted by *open circles*. Amino acids selected for mutagenesis are numbered. **c**, Disorder plots and charge heatmaps are shown for the CRD (species and details are as described in panel *a*). **d**, Scatter plots of isoelectric point, disorder score, disorder %, and disorder length for CBX5, CRD, KMT5C, and MeCP2 across 20 representative mammalian species (Extended Data Table 1). **e**, Partition images (16-color intensity map) for the CRD from *Hs, Mm, Cp*, and *Bb* indicate that all effectively partition to chromocenters. **f**, Partition coefficient graph for *Hs* (*n=39*), *Mm* (*n=30*), *Cp* (*n=23*), and *Bb* (*n=26*) CRDs demonstrate high chromocenter partitioning. Full-length KMT5C is shown for reference (*faded*). **g**, FRAP analyses of *Hs, Mm, Cp*, and *Bb* CRD domains (Extended Data Movies 6-9) support intra-chromocenter mobility with reduced nucleoplasmic exchange (*n=30*).

We next examined the role of conserved residues in the CRD, focusing on C^362^C^366^, H^357^H^365^, and W^359^W^390^W^392^Y^396^ (Fig. 2b), because they resembled features found in chromatin reader modules that jointly recognize DNA and histones.^18^ Mutation of individual motifs caused elevated exchange in FRAP assays (Fig. 3a, Extended Data Videos 10-12), which was accompanied by increased nucleoplasmic abundance (Fig. 3b) and decreased partitioning (Fig. 3c, d). Combining mutants (C^366^W^390^W^392^) abrogated chromocenter localization and led to rapid mobility (Fig. 3a-d, Extended Data Video 13). These findings established that the CRD has evolved multiple determinants that cooperate to confer localization and limit its exit from individual chromocenters, while still supporting a liquid-like state. Non-membranous organelles that form by phase separation are typically spherical, reflecting a reduction in their surface area,^19^ which prompted us to evaluate this parameter in CRD mutants (Fig. 3e). Importantly, each of the mutants caused a reduction in sphericity that correlated with their partitioning and dynamic behavior (Fig. 3a-d), with C^362^C^366^ being most severe and W^359^W^390^W^392^Y^396^ the least. The extent mutants perturbed CRD activity therefore reflected the degree to which they reduced the valency or interaction affinity, a finding that is consistent with the behavior of other phase separated proteins.^13^ We extended this analysis to include KMT5C, CBX5, MeCP2, and the remaining CRDs. Strikingly, KMT5C supported a significantly higher degree of sphericity than either CBX5 or MeCP2 (Extended Data Fig. 1a), indicating it was more effective at generating surface tension at the boundary between the chromocenter and nucleoplasm. Moreover, all CRDs shared this property (Extended Data Fig. 1b), further highlighting that this domain is the key determinant of the biophysical characteristics of KMT5C.

**Figure 3.**
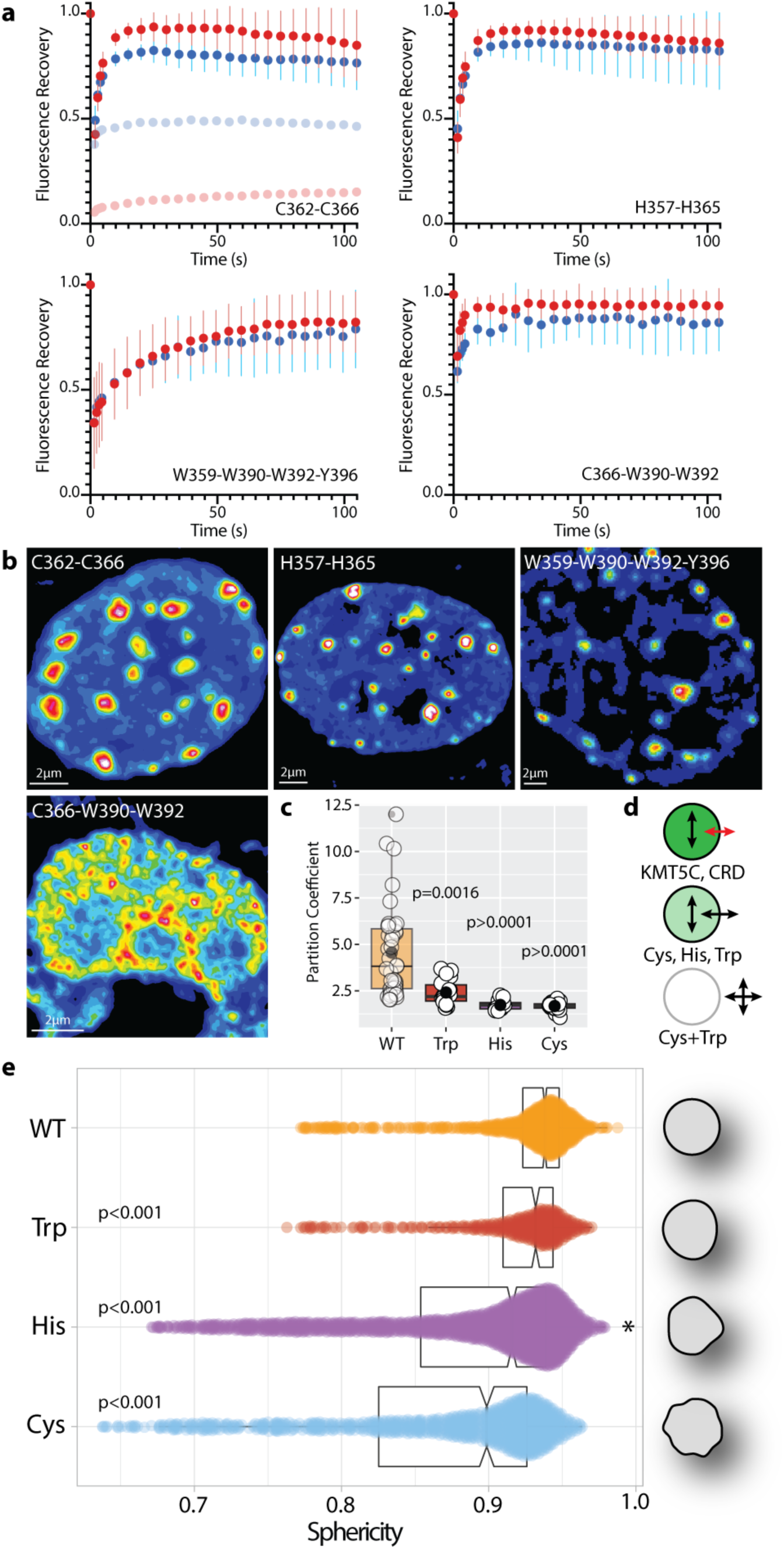
The CRD comprises multiple determinants that cooperate to drive heterochromatin phase separation. **a**, FRAP analysis of CRD mutants that target conserved sequence features (C^362^C^366^, H^357^H^365^, and W^359^W^390^W^392^Y^396^) or combinations thereof (C^366^W^390^W^392^) (*n=30*). Wild-type KMT5C (*faded*) is shown in the upper left panel for reference (Extended Data movies 10-13). While the recovery profiles are similar for the C^362^C^366^ and H^357^H^365^ mutants, they differed for the W^359^W^390^W^392^Y^396^ and mutants, suggesting they affect distinct interactions. **b**, Partition images (16-color intensity map) for CRD mutants. **c**, Partition coefficients for CRD mutants (WT (*n=39*), Trp (*n=18*), His (*n=17*), and Cys (*n=20*)). **d**, Schematic summary illustrates that mutants continue to localize to the chromocenter, but now readily exchange with the cytoplasmic pool. **e**, Sphericity analysis of the wild-type (*n=990*) and mutant (Trp (*n=437*), His (*n=1834*), and Cys (*n=1054*) CRDs (see Methods).

Phase separation by CBX5 is abrogated in *Suv39h1/2* null cells or by mutation that disrupts H3K9me3 recognition,^1^ underscoring the importance of this interaction in seeding liquid demixing. We found the dynamic behavior of KMT5C exhibited the same dependency, reflected by its increased mobility and dispersal in cells lacking SUV39H1/2 (Fig. 4a, Extended Data Videos 14, 15). MECP2, however, displayed more efficient localization and slightly reduced mobility (Extended Data Fig. 2), indicating the chromocenter can support distinct phase separated assemblies depending on chromatin context. Another facet of phase separation involves its reversibility in response to cellular queues.^13^ To this end, we evaluated the histone deacetylase inhibitor Trichostatin A (TSA) because it was known to cause CBX5 displacement from pericentromeric heterochromatin and increase its mobility.^20^ KMT5C, however, largely retained chromocenter localization, but with elevated nucleoplasmic exchange (Fig. 4b, Extended Data Video 16), indicating hyperacetylation differentially affects KMT5C and CBX5 demixing in cells. Nevertheless, this modest KMT5C release was associated with a marked accumulation of H4K20me3 outside of chromocenters (Fig. 4b), establishing that retention is essential to the spatial regulation of its enzymatic activity, which otherwise acts promiscuously. Alteration of KMT5C dynamics was also apparent upon induction of DNA damage within chromocenters by laser microirradiation, which is known to induce heterochromatin decompaction and changes in histone post-translational modifications.^21^ Specifically, bisection of chromocenters using laser microirradiation caused loss of KMT5C from the damaged area, creating two lobes that preserved the mobility and retention behavior of the original domain (Fig. 4c, Extended Data Videos 17, 18). Unlike the global change observed with TSA treatment, this reflected a locally confined and precise dissolution in response to underlying chromatin state changes over a timescale of seconds. Again, the behavior of KMT5C was markedly different than either CBX5 or MeCP2, which showed residual localization in the damage zone and rapid mobility (Fig. 4c, d, and Extended Data Videos 19-22). Collectively, these findings indicate that phase separation can tune constitutive heterochromatin protein content and dynamics in response to changes in chromatin state and environmental stimuli.

**Figure 4.**
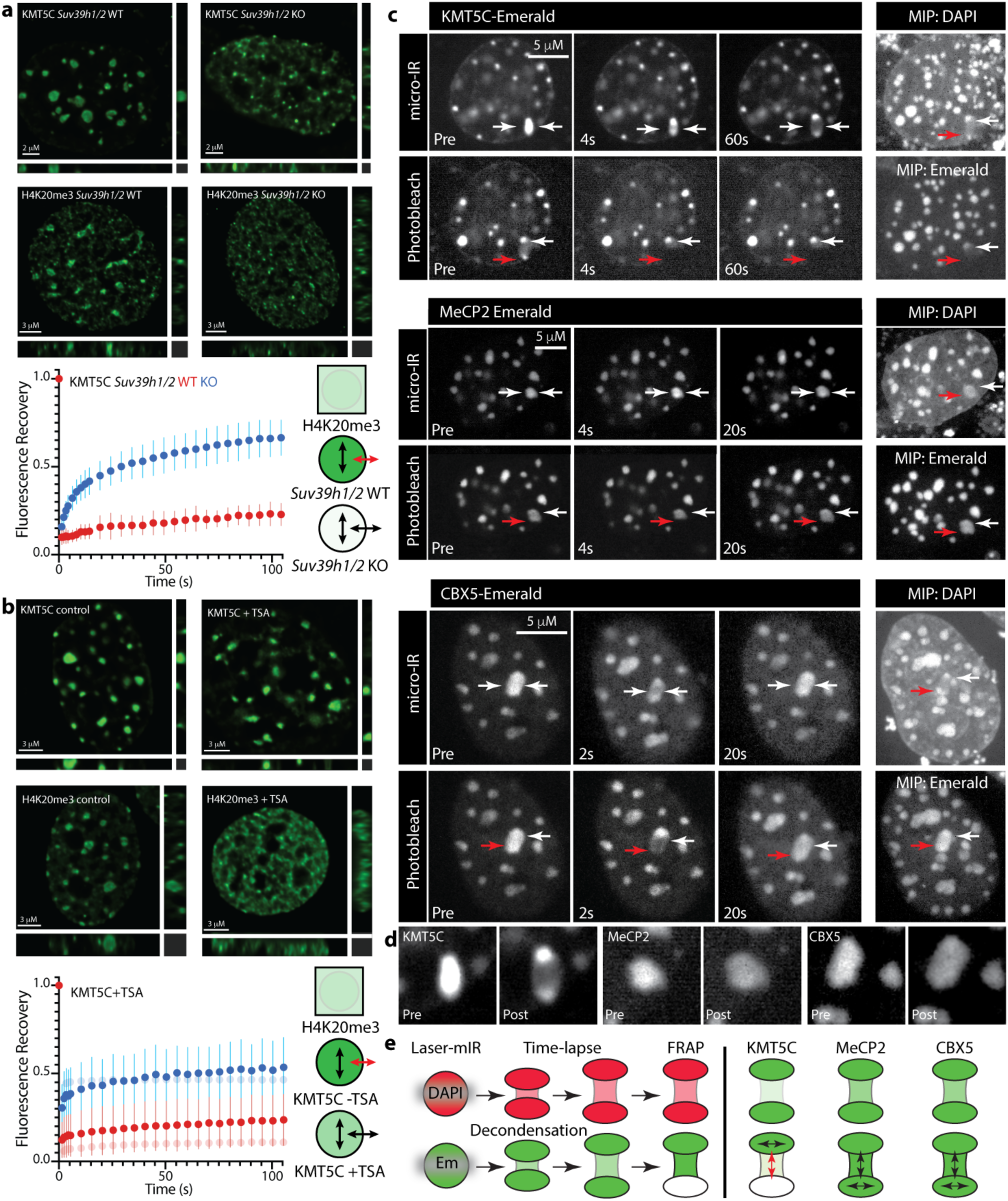
KMT5C phase separation is rapidly reversible and responsive to underlying chromatin state. **a**, KMT5C localization and mobility in *Suv39h1/2* knockout cells. *Top row*, KMT5C localization is dependent on H3K9me3 placement by SUV39H1/2^35^ which leads to redistribution of H4K20me3 (*middle row*) and marked in increase in KMT5C mobility in *Suv39h1/2* knockout cells when assessed by FRAP (*bottom row* and Extended Data Videos 14, 15) (*n=30*). **b**, The histone deacetylase inhibitor trichostatin A (TSA) alters KMT5C localization and mobility, and leads to H4K20me3 redistribution (treatment was for 16-24hrs at 100nM). *Upper row*, KMT5C localization is only moderately affected by TSA treatment, but H4K20me3 undergoes dramatic redistribution (*middle row*) that coincides with increased KMT5C nucleoplasmic exchange in FRAP (*bottom row* and Extended Data Video 16) (*n=30*). In *a* and *b*, the schematic indicates H4K20me3 is no longer confined to the chromocenter under either condition, but that *Suv39h1/2* knockout had a larger effect on KMT5C localization, despite similar increases in mobility with TSA treatment. **c**, Laser micro-irradiation of KMT5C, MeCP2, and CBX5 differentially modulates their phase separation (Extended Data Videos 17-22). For each protein, the *top row* represents a time series that includes pre-damage (Pre), immediately following damage (4s), and at 60s (area targeted by micro-IR is indicated by facing *arrows*). The *bottom row* depicts a FRAP time series where a portion of the damaged chromocenter (*arrows* in Pre-bleach) is photobleached (*red arrow*) and recovery is monitored at 4s and either 60s (KMT5C) or 20s (MeCP2 and CBX5). Maximum image projections (MIP) are included to highlight the area of decondensed heterochromatin following damage induction and the location of the bleaching zone for mEmerald fusions of KMT5C, MeCP2, and CBX5. **d**, Comparison of pre and post irradiation images for KMT5C, MeCP2 and CBX5. While MeCP2 and CBX5 behaved similarly and underwent only transient or no decrease in intensity over the damaged area, KMT5C rapidly exited and accumulated in the two non-damaged lobes. **e**, Schematic summary of laser microirradiation and FRAP analyses. The *left* panel summarizes the experimental strategy, which involves using a laser to induce chromocenter DNA damage and then characterize the behavior of the exogenously expressed fusion protein, as well as its capacity to recover fluorescence following photobleaching. The DAPI series was used to monitor heterochromatin decompaction following damage, while the response of the mEmerald fusion protein was visualized over time at which point the lower portion of chromocenter was bleached in order to determine protein mobility. In both contexts, KMT5C behaved markedly different than MeCP2 and CBX5. It neither persisted in the damaged area nor exhibited fluorescence recovery.

The CRD provides a minimalist model to decipher how phase separation can support spatial confinement while at the same time maintaining liquid-like behavior. Although lacking many of the typical sequence features of phase separating proteins,^3,13^ the CRD met the criterion of multivalency. Nevertheless, while this normally controls the composition and biophysical properties of phase separated assemblies via low affinity interactions that allow proteins to exchange with their surroundings,^3^ the CRD underwent very limited exchange. In this regard, CRD mutants resembled full-length CBX5 in mobility and partitioning, indicating multiple determinants act in a highly synergistic manner to self-reinforce chromocenter retention. This behavior was independently supported by the response of KMT5C to inhibition of HDACs (Fig. 4), which partially diminished retention. For *Drosophila* HP1a, H3K9me3 recognition is driven by cation-*π* interactions involving a triad of aromatic residues (Y^24^, W^45^, and Y^48^) within the chromodomain,^15^ and is necessary for heterochromatin phase separation.^2^ A key difference in the CRD is the presence of five aromatic residues (W^359^, W^384^, W^390^, W^392^, and Y^396^) and the predominance of tryptophan, which confers the strongest cation-*π* interactions^22^ and supports more stable binding of trimethyllysine.^23^ Together with multivalency and the capacity for *π*-*π* interactions,^24^ the CRD appears to have evolved unique determinants that are optimized to confine KMT5C to constitutive heterochromatin. Moreover, when compared to other phase separating proteins, this activity has been consolidated into a limited number of sequence features that are sufficient to reduce chromocenter surface area and enable exquisite control of protein localization in the aqueous phase.

Phase separation can be described in phase diagrams where changes in protein concentration and interaction strength give rise to assemblies with distinct material properties,^19^ including gels, glassy solids, and pathological aggregates.^13^ The latter are states where molecules are immobile, retain their relative positions to each other, and do not exchange with their surroundings.^19^ This is clearly not what we have described for KMT5C, which remained in a liquid state despite limited exchange that initially suggested it was immobile.^12^ Although in principle high partitioning to a phase separated compartment has the potential to drive retention,^3^ the behavior of MeCP2 indicates that this feature alone is not sufficient, but also requires a high energy barrier to exit. For KMT5C, this combination allows it to be effectively biocontained within a phase separated ‘organelle’ and affords tight spatial control of H4K20me3 catalysis. This is notable given it constitutes a minority of the methylated H4K20 pool^25^ and is primarily associated with satellite and other distinct repeats.^26^ This control is lost when KMT5C is no longer constrained in demixed assemblies (Fig. 4). Moreover, the opposing behaviors of KMT5C and MeCP2 indicated that change of a single epigenetic feature can dramatically reprogram the phase separated state with regard to protein composition and dynamics. Of particular relevance to this finding, MeCP2 loss in a mouse model of Rett syndrome leads to H4K20me3 gain in the chromocenter, supporting an antagonistic relationship with KMT5C *in vivo* at endogenous protein levels.^27^ This principle is also supported by an altered chromocenter proteome in *Suv39h1/2* knockout cells^28^ and the established plasticity of constitutive heterochromatin in development and disease.^29^ The existence of these distinct phase separated states therefore provides a conceptual framework to understand the drivers of normal and pathogenic chromocenter homeostasis. By considering chromatin as a multivalent scaffold, we can decipher how changes in epigenetic features and the mutational status of resident proteins shift the composition and function of phase separated assemblies.

## Methods

### Cell Culture and transfection

Cells were cultured at 37°C and 5% CO_2_ in a humidified incubator. All cell lines were grown in DMEM containing 10% FBS. D5 (*Suv39h1/2* knockout) and W8 (*Suv39h1/2* wild-type) mouse embryonic fibroblast cells lines^8^ were obtained from Dr. Thomas Jenuwein. All other analyses were carried out using the mouse NMuMG breast cell line. Cells were transfected by lipofection using Effectene (Qiagen) 1 day prior to experiments. Expression plasmids were synthesized (www.biomatik.com), obtained from the Addgene repository (www.addgene.org), or previously described^30^ (sequences provided in supplemental material).

### Live-cell imaging

Live cell imaging was carried out using Zeiss Axiovert 200M inverted microscopes attached to either an LSM510 NLO laser scanning system with a 25 mW argon laser line, a Zeiss LSM 770 confocal microscope attached to an Axio Observer Z3 equipped with 405, 488, 561, and 633 nm diode lasers, or a PerkinElmer Ultraview spinning-disk confocal microscope equipped with 405, 488, and 561 nm diode lasers and a FRAP-unit. For all platforms, a 40 x 1.3 NA oil immersion lens was used. Long-term live-cell observations were conducted on the spinning disk microscope at 37°C in a humidified atmosphere containing 5% CO_2_. In cases were Z-stacks were acquired, spacing was set at 400 nm. Fluorescence recovery after photobleaching was performed on transiently transfected cells using the 488 nm solid state (spinning disk confocal) or 488 nm argon laser line (LSM 510). Circular (chromocenter) or linear (nucleoplasm) regions were demarcated and subsequently bleached by intense light from the 488 nm laser. Fluorescence recovery of the bleached regions was quantified over multiple time scales (seconds to minutes). FRAP data was extracted using Zeiss LSM 5 Zen or ImageJ software by measuring fluorescence intensity of the background, the whole nucleus and the bleached area in each of the recorded time-lapse pictures for a minimum of 30 cells. Normalized relative intensity (including standard deviation) was calculated in Microsoft Excel and plotted using Graphpad Prism software. Laser microirradiation experiments were performed using the spinning disk microscope using the 100 × 1.4 NA objective lens. Cells were grown in MatTek 35 mm glass bottom dishes and were sensitized with 1 µg/µl Hoechst 20 min prior to the experiment. After calibration of the photokinesis device, a thin horizontal line representing the region to be microirradiated was placed so that it divided the chromocenter into approximately two equal parts. Laser microirradiation was carried out by using 20% power of the 405 nm solid state laser and 10 iterations. Images were acquired at defined time intervals using laser and filter settings for GFP imaging. Subsequent photobleaching of the non-microirradiated portion of chromocenters was done using 10 iterations at 100% power of the 488 nm solid state laser.

### Immunofluorescence

Cells grown on adherent coverslips were fixed in 4% paraformaldehyde in PBS for 10 min, permeabilized in 0.5 % Triton X-100 in PBS for 5min and then incubated with primary antibodies diluted in PBS. After 30 min at room temperature, the cells were washed once with 0.1 % Triton X-100 in PBS for 1 min then rinsed 3 times with PBS prior to addition of secondary antibodies diluted in PBS. Cells were incubated for 30 min at room temperature, washed with 0.1 % Triton X-100 in PBS for 1 min and rinsed three times with PBS. Coverslips were mounted onto microscope slides with in-house made polyvinyl alcohol mounting media containing 1 µg/mL 4′,6-diamidino-2-phenylindole (DAPI). Z-Stacks were obtained on a Zeiss Imager.Z1 equipped with a Photometrics Prime BSI camera and Metamorph software version 7.10.2.240 (Molecular Devices, Sunnyvale, CA) using a Zeiss 63 × 1.3 NA oil lens. Step size used was 0.2 µm. Primary antibodies and dilutions were as follows: *α*-H4K20me3 (Active Motif 39672), 1:500; and *α*-H3K9me3 (Active Motif 3916). Secondary antibodies and dilutions were as follows: goat *α*-mouse Alexa-488, 1:500; goat *α*-rabbit Alexa-488 1:500; and goat *α*-mouse Cy3, 1:500.

### Image analysis

Images were deconvolved with Huygens Professional version 19.04 (Scientific Volume Imaging, http://svi.nl). Z-stacks were imported into Imaris 9.3 (Oxford Instruments) and cropped to generate 3D images of single nuclei. The Imaris *surfaces* function was used to encapsulate all chromocentres with a shell and the *statistics* function was used to measure the volume, sphericity, and total number of chromocenters for output to a Microsoft Excel spreadsheet. Graphical representation of the data was prepared using the PlotsOfDifferences server (https://huygens.science.uva.nl/PlotsOfDifferences/).^31^ For partition coefficients, line scans through nuclei of undeconvolved 3D images were recorded using ImageJ. For each cell, the partition coefficient was calculated by subtracting the background level from the maxima of the brightest chromocenter and dividing it by the background corrected fluorescence intensity of the nucleoplasm. For each protein analyzed, measurements were taken from n>10 nuclei.

### Statistical analysis

For partition coefficients, significance was evaluated using the Kruskal-Wallis one-way analysis test and individual comparisons between proteins were done using the Wilcoxon rank sum test. Statistical significance of sphericity data was evaluated using the embedded stats function within PlotsOfDifferences, which calculates p-values using a randomization test.^31^

## Extended Data

### Videos

Video 1; KMT5C FRAP

Video 2; MeCP2 FRAP

Video 3; CBX5 FRAP

Video 4; Partial KMT5C bleach (doughnut)

Video 5; Partial KMT5C bleach (bridge)

Video 6; *Homo sapiens* CRD FRAP

Video 7; *Mus musculus* CRD FRAP

Video 8; *Cavia porcellus* CRD FRAP

Video 9; *Bubalus bubalis* CRD FRAP

Video 10; CRDC^362^C^366^ FRAP

Video 11; CRDH^357^H^365^ FRAP

Video 12; CRDW^359^W^390^W^392^Y^396^ FRAP

Video 13; CRDC^366^W^390^W^392^ FRAP

Video 14; KMT5C in D5; *Suv39h1/2* knockout FRAP

Video 15; KMT5C in W8; *Suv39h1/2* wild-type FRAP

Video 16; KMT5C with TSA FRAP (no TSA control; Video 1)

Video 17; KMT5C laser microirradiation

Video 18, KMT5C laser microirradiation + FRAP

Video 19; MeCP2 laser microirradiation

Video 20, MeCP2 laser microirradiation + FRAP

Video 21; CBX5 laser microirradiation

Video 22, CBX5 laser microirradiation + FRAP

## Figures and Tables

Table 1; Spreadsheet 1; PCH PONDR excel; additional species disorder and physicochemical data for KMT5C, MeCP2, CBX5, and CRD

Figure 1; Sphericity data for (a) KMT5C, MeCP2, CBX5, (b) *Hs, Cp, Bb*, and *Cp* CRDs, and (c) wild-type and mutant CRDs.

Figure 2; MeCP2 (a) localization and (b) mobility in D5 (*Suv39h1/2* knockout) and W8 (*Suv39h1/2* wild-type) cells

## Acknowledgements

The authors acknowledge funding support from the Canadian Breast Cancer Foundation (D.A.U., grant no. 300073), Cancer Research Society (D.A.U. and M.J.H., grant no. CRSDI 2018 OG 23446), and Canadian Institutes of Health Research (M.J.H., grant no. PJT-148753). The authors thank Dr. Xuejun Sun and Gerry Barron of the Cross Cancer Institute Cell Imaging Facility for support.

## Author Contributions

D.A.U. and M.J.H. conceived the project. D.A.U., M.J.H, and H.S. designed all experiments and interpreted data. K.M. made the initial observation of KMT5C mobility and performed FRAP and fluorescence imaging experiments. H.S. performed live-cell imaging, laser micro-irradiation, and partition coefficient analyses. D.A.U. carried out sequence analysis and expression plasmid design. D.A.U. wrote the manuscript with contributions from M.J.H. and H.S. D.A.U. prepared figures with contributions from H.S. and K.M.

## Containing data deposition statement

N/A

## Extended Data Figures and Legends

**Figure 1.**
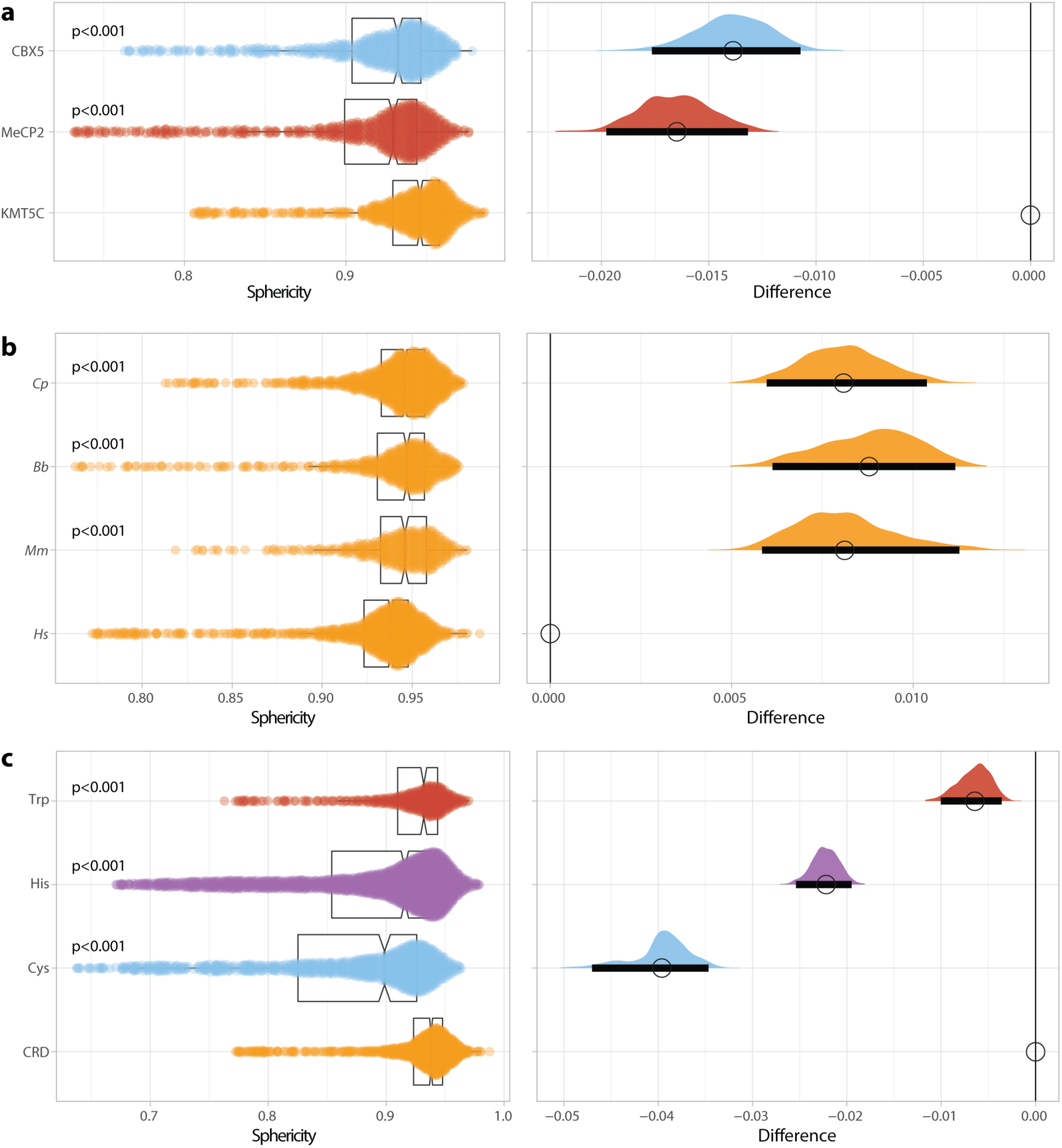
Chromocenter sphericity analysis of KMT5, MeCP2, CBX5, and all CRD derivatives. **a**, Comparison of chromocenter sphericity of full-length KMT5C (*n=775*), MeCP2 (*n=641*), and CBX5 (*n=644*) indicates KMT5C supports significantly higher sphericity. **b**, Comparison of chromocenter sphericity generated by CRDs from *Homo sapiens* (*Hs*; *n=990*), *Mus musculus* (*Mm*; *n=421*), *Bubalus bubalis* (*Bb*; *n=574*), and *Cavia porcellus* (*Cp*; *n=982*). All CRDs support efficient sphericity, although this is significantly higher for *Mm, Bb*, and *Cp* when compared to *Hs*, which also exhibited reduced chromocenter portioning (*n.s*.) (manuscript Fig. 2f) 2. Nevertheless, the human CRD supports significantly higher sphericity than MeCP2 (p<0.001) and CBX5 (p<0.001), or CRD mutants (*panel c*). **c**, Comparison of sphericity for wild-type (*Hs*) and mutant CRDs (as in manuscript Fig.3e, with inclusion of difference plot). All proteins were assessed in murine NMuMG breast cancer cells (see Methods).

**Figure 2.**
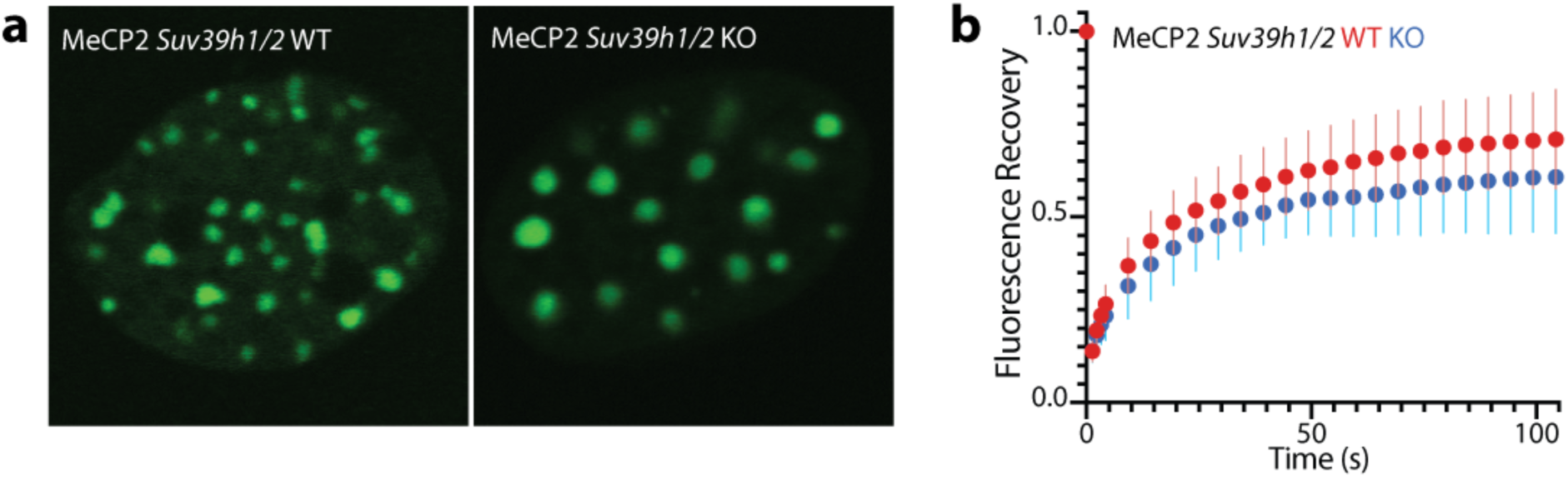
Localization and FRAP of MeCP2 in *Suv39h1/2* wild-type and knockout cells. , MeCP2-GFP displays efficient chromocenter localization in both *Suv39h1/2* wild-type and knockout mouse embryonic fibroblasts. **b**, FRAP analysis of MeCP2-GFP reveals a decrease in mobility in *Suv39h1/2* knockout cells.

